# Growth of Gram-negative uropathogens in human urine under hypoxic conditions produces clinically relevant metabolomic profiles

**DOI:** 10.1101/2025.05.07.652693

**Authors:** Mohamed Eladawy, Anne L. McCartney, A. Christopher Garner, Lesley Hoyles

**Affiliations:** Department of Biosciences, School of Science and Technology, Nottingham Trent University, Nottingham, UK; Department of Microbiology and Immunology, Faculty of Pharmacy, Mansoura University, Mansoura, Egypt; Department of Chemistry, School of Science and Technology, Nottingham Trent University, Nottingham, UK

**Keywords:** urinary tract infections, bladder, *Escherichia coli*, *Klebsiella pneumoniae*

## Abstract

**Background:** Much work on uropathogens is done in rich laboratory media that do not reflect the nutrient availability of urine. Also, the environment of the bladder is microaerobic (∼5 % O_2_) but routine laboratory work with uropathogens is done under aerobic conditions.

**Aims:** To understand the influence of oxygen conditions (aerobic, microaerobic) and physiologically relevant growth substrates (artificial urine, human urine) on the ability of bacteria isolated from catheter-associated urinary tract infections (CAUTIs) to form biofilms, and to begin to define the CAUTI bacterial metabolome.

**Methods:** Biofilm assays were conducted in rich lab medium (tryptone soy broth supplemented with glucose), and commercially available artificial and pooled human urine for 29 well-characterized uropathogens representing five different genera of Gram-negative bacteria. Spent media were collected and analysed using 400 MHz ^1^H-NMR, to quantify major metabolites produced by uropathogens in artificial urine and human urine.

**Findings:** There was a significant decrease in biofilm formation for all uropathogens when grown in physiologically relevant growth substrates. Untargeted metabolomic analyses of artificial and human urine showed the artificial urine used in this study did not recapitulate the complexity of major metabolites present in human urine. Further analyses of spent human urine highlighted significant increases in acetate production by the uropathogens when they were grown under microaerobic compared with aerobic conditions.

**Conclusions:** Growth of uropathogenic *Enterobacteriaceae* under physiologically relevant conditions (i.e. human urine, 5 % O_2_) generates data more relevant to clinical disease and is an important consideration for future work on bacteria causing urinary tract infections.

## INTRODUCTION

Urinary tract infections (UTIs) occur in both community and healthcare settings, ranging from uncomplicated to complicated infections. Over half of all women experience at least one UTI during their lives, with over a quarter experiencing a recurrence 3–6 months after their initial infection [1]. The likelihood of experiencing a recurrent UTI increases with age, with postmenopausal women twofold more susceptible than premenopausal women [1]. Bladder-associated UTIs, such as cystitis, are more common in women than men [2]. UTIs confer a significant economic burden on healthcare systems, representing approximately 40 % of healthcare-associated infections [2,3]. In 2019, there were ∼405 million cases of UTIs, with ∼237,000 deaths and ∼5.2 million disability-associated lost days linked to these infections [4]. Deaths associated with UTIs increased more than twofold between 1990 and 2019, accompanied by an increasing age-standardized mortality rate over time [4].

Complicated UTIs are often associated with catheterization, with uropathogens able to bind to the catheter in the bladder and multiply under the protection of the biofilm they produce [2]. It is estimated that ∼2 % of patients hospitalized for more than 48 hours develop a nosocomial UTI, and that ∼65 % of catheter-associated urinary tract infections (CAUTIs) may be preventable [5]. If left untreated, CAUTIs can progress to pyelonephritis and bacteraemia.

Gram-negative bacteria cause most healthcare-associated UTIs [2]. Uropathogenic *Escherichia coli* (UPEC) causes most infections (∼65 %) in western clinical settings, followed by *Klebsiella pneumoniae* (∼8 %), *Proteus mirabilis* (∼3 %) and *Pseudomonas aeruginosa* (∼3 %) [2]. Diagnosis of a UTI is made by culturing urine samples under aerobic conditions and obtaining a bacterial count ≥10^5^ cfu/ml, though patients with symptomatic UTIs can have bacterial counts 10^3^ to <10^5^ cfu/ml [6,7]. Antimicrobial-resistant UTIs represent the fourth most prevalent infectious syndrome associated with global deaths directly linked to antimicrobial resistance [4].

While the recent ENACT report has addressed behavioural changes that can be made in hospital settings to reduce incidences of CAUTIs [8], the microbiological processes associated with these infections remain poorly studied. Much of the microbiological research undertaken in this area relies on growth of bacteria in rich laboratory media and in the presence of air (21 % O_2_). However, urine is a nutrient-limited substrate, and the bladder represents a hypoxic environment (0.06–6.8 % O_2_) [9–12]. There are many confounders affecting the hypoxic environment of the bladder, including factors such as systemic metabolic health and oxygenation, local diseases of the urinary tract and/or kidneys, renal blood flow, urine flow, UTIs, diuretics and chronic kidney disease [13]. UTIs reduce the availability of dissolved oxygen in urine [12].

Metabolite profiling of urine samples using ^1^H-nuclear magnetic resonance spectroscopy (NMR) has been proposed as a means of rapid diagnosis of UTIs, with elevated levels of acetate considered a biomarker of these infections [14–17]. Elevated levels of trimethylamine (TMA), along with acetate, have been considered a biomarker for UPEC-associated UTIs [15,18]. Both acetate (trace to 100 µM/mM creatinine) and TMA (trace to 15 µM/mM creatinine) are gut-microbiome-associated metabolites found in human urine (HU), normally produced through microbial breakdown of dietary substrates in the gastrointestinal tract and contributing to the endogenous exposome [19–22]. Other gut-microbiome-associated metabolites routinely detected in HU by NMR include trimethylamine *N*-oxide (TMAO; 36–202 µM/mM creatinine), dimethylamine (DMA; 31–47 µM/mM creatinine) and hippurate (20–770 µM/mM creatinine) [22]. If the levels of these metabolites are increased in the urine of patients with UTIs, it is possible that uropathogens are producing acetate, DMA and/or TMA from components of HU. Hippurate and TMAO are host–microbiota co-metabolites: conjugation of microbially produced benzoate with glycine leads to hippurate production [23], while TMAO is produced from TMA by hepatic flavin monooxygenases [20,24]. As such, microbes in urine should not produce hippurate or TMAO from constituents of urine.

Systems-level work in metabolic diseases has shown that host–microbiota interactions and microbially produced metabolites influence disease progression [20,25,26]. There is a need to consider UTIs in a similarly holistic manner. A better understanding of how bacteria form biofilms in the urinary tract and metabolize components of HU will ultimately lead to greater understanding of infection initiation and progression, and to identification of novel therapeutic agents to prevent or reduce severity of UTIs. As such, the aims of our study were to determine the biofilm-forming abilities of CAUTI bacteria under physiologically relevant conditions [i.e. in artificial urine (AU) or HU under hypoxia], and to determine which metabolites CAUTI-associated bacteria produce under these conditions.

## METHODS

### Strains included in this study

All uropathogens (*n* = 29) used in this study (**Table 1**), including details of their isolation from CAUTIs and their characterization, have been described elsewhere [27].

**Table 1.**
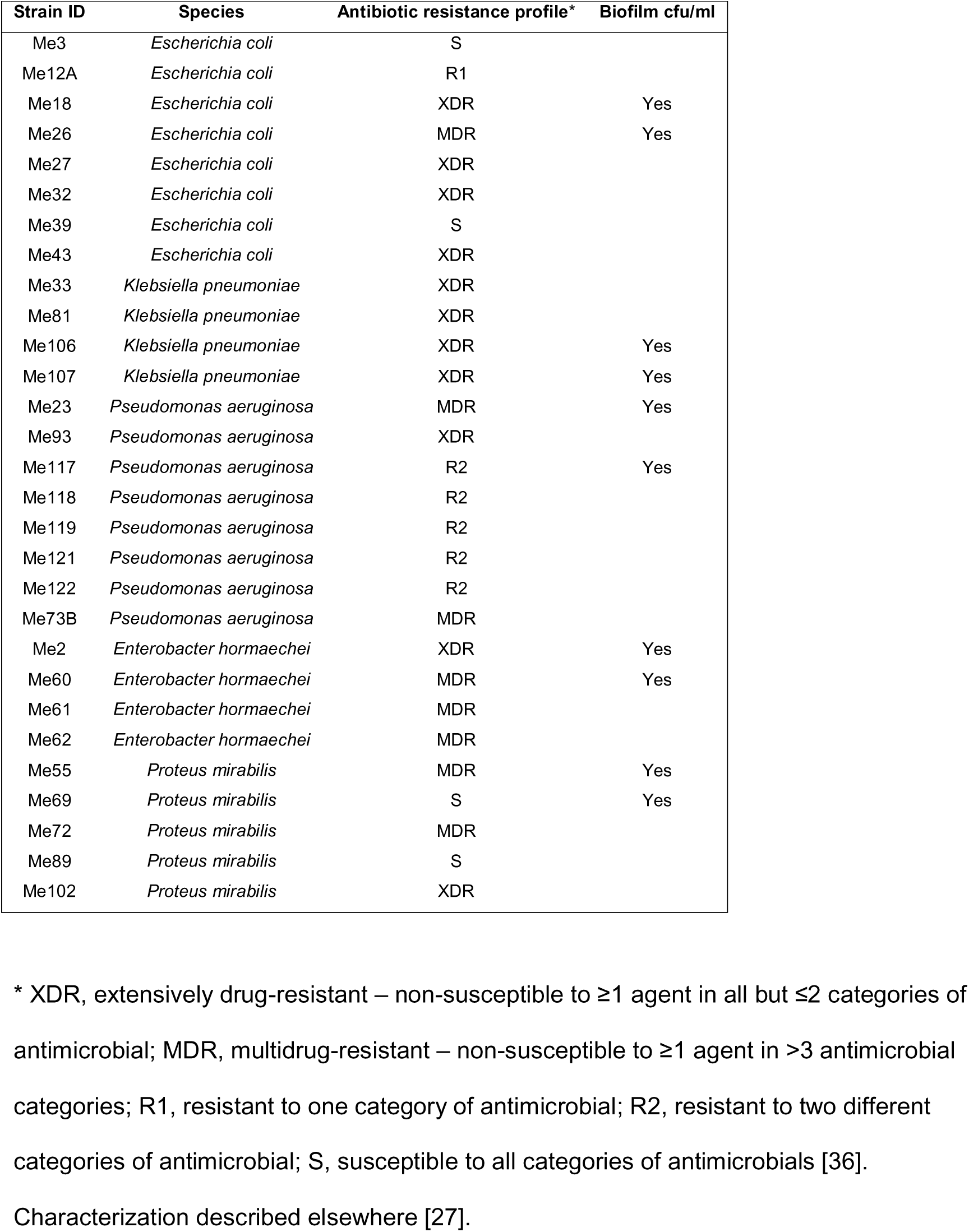
Details of strains included in this study.

| Strain ID | Species | Antibiotic resistance profile* | Biofilm cfu/ml |
| --- | --- | --- | --- |
| Me3 | *Escherichia coli* | S |  |
| Me12A | *Escherichia coli* | R1 |  |
| Me18 | *Escherichia coli* | XDR | Yes |
| Me26 | *Escherichia coli* | MDR | Yes |
| Me27 | *Escherichia coli* | XDR |  |
| Me32 | *Escherichia coli* | XDR |  |
| Me39 | *Escherichia coli* | S |  |
| Me43 | *Escherichia coli* | XDR |  |
| Me33 | *Klebsiella pneumoniae* | XDR |  |
| Me81 | *Klebsiella pneumoniae* | XDR |  |
| Me106 | *Klebsiella pneumoniae* | XDR | Yes |
| Me107 | *Klebsiella pneumoniae* | XDR | Yes |
| Me23 | *Pseudomonas aeruginosa* | MDR | Yes |
| Me93 | *Pseudomonas aeruginosa* | XDR |  |
| Me117 | *Pseudomonas aeruginosa* | R2 | Yes |
| Me118 | *Pseudomonas aeruginosa* | R2 |  |
| Me119 | *Pseudomonas aeruginosa* | R2 |  |
| Me121 | *Pseudomonas aeruginosa* | R2 |  |
| Me122 | *Pseudomonas aeruginosa* | R2 |  |
| Me73B | *Pseudomonas aeruginosa* | MDR |  |
| Me2 | *Enterobacter hormaechei* | XDR | Yes |
| Me60 | *Enterobacter hormaechei* | MDR | Yes |
| Me61 | *Enterobacter hormaechei* | MDR |  |
| Me62 | *Enterobacter hormaechei* | MDR |  |
| Me55 | *Proteus mirabilis* | MDR | Yes |
| Me69 | *Proteus mirabilis* | S | Yes |
| Me72 | *Proteus mirabilis* | MDR |  |
| Me89 | *Proteus mirabilis* | S |  |
| Me102 | *Proteus mirabilis* | XDR |  |

\* XDR, extensively drug-resistant – non-susceptible to $\geq 1$ agent in all but $\leq 2$ categories of antimicrobial; MDR, multidrug-resistant – non-susceptible to $\geq 1$ agent in $>3$ antimicrobial categories; R1, resistant to one category of antimicrobial; R2, resistant to two different categories of antimicrobial; S, susceptible to all categories of antimicrobials [36]. Characterization described elsewhere [27].

### Biofilm assay and collection of spent media

Biofilm assays were performed as described previously [28], but using three different growth media – tryptone soya broth (Oxoid Ltd) plus 1 % glucose (TSB; [28]), non-stabilized AU (LC Tech; catalogue no. 1700-0018; lot no. 2303006) and pooled, filtered HU (BioIVT; catalogue no. HUMANURINE-104655; lot no. HMN1084681) – and two different atmospheric conditions: aerobic and microaerobic (Don Whitley DG250 workstation; 5 % O_2_, 10 % CO_2_, 85 % N_2_).

Initially, a colony of each strain was inoculated into LB broth, and incubated at 37 °C overnight. After incubation, 1 ml from each overnight culture was centrifuged at 13000 rpm for 5 min, and the supernatant discarded. Pellets were washed with phosphate-buffered saline (PBS; pH 7.4; Oxoid Ltd) twice to remove any traces of spent medium. The pellets were resuspended in 1 ml of PBS. The optical density (OD) of cell suspensions was adjusted to OD_600_ 0.2 (∼10^8^ cfu/ml). For each strain, 10 μl of the OD-adjusted culture was inoculated into three wells of a 96-well, flat-bottomed polystyrene microtitre plate containing 90 μl of TSB, AU or HU. Negative controls containing sterile TSB, AU or HU were included in each experiment. The 96-well plates were incubated aerobically or microaerobically, without shaking, at 37 °C for 24 h. Then, the liquid culture was carefully removed from each well. For each strain, HU- or AU-grown cultures were pooled, filtered through 0.2 μm sterile syringe filters (Minisart, Sartorius) and stored at −80 °C in sterile microcentrifuge tubes. The adhered biofilm left at the bottom of each well was carefully washed three times with 200 μl of PBS to remove any non-adherent cells.

#### Crystal violet assay

Adherent cells were fixed by addition of 150 μl of absolute methanol (Fisher Scientific, catalogue no. 11480520) for 15 min. Afterwards, the methanol was aspirated, and the adhered biofilm was stained by addition of 1 % (w/v) crystal violet solution (Fisher Scientific, catalogue no. 12916287; 150 μl per well) and the plate was left for 20 min. The plate was then carefully rinsed with distilled water three times and kept inverted to air dry the wells. The stained biofilm was solubilized by adding 33 % (v/v) glacial acetic acid (Fisher Scientific, catalogue no. 10365020; 150 μl per well). After solubilization, the *A*_540_ was measured using an Elx808™ absorbance microplate reader.

### Determination of colony-forming units (cfu) of bacteria in biofilms

Biofilm assays were set up and non-adherent cells removed as described above. For each set of technical replicates (*n* = 3) within a biological replicate for AU or HU, after aerobic or microaerobic incubation for 24 h 200 μl of sterile PBS was added to each well and the biofilms were scraped from the wells using the end of an inoculation loop. The samples were then pooled (total volume 600 μl), and 400 μl of PBS was added to the pooled samples, which were vortexed before being used to make a dilution series that was plated onto nutrient agar (Oxoid Ltd) to determine the number of cfu/ml of sample.

### NMR

Buffer prepared in deuterated H_2_O [0.2 M phosphate (Na_2_HPO_4_, NaH_2_PO_4_) buffer, pH 7.4; 0.2 % NaN_3_, 1 mM 3-(trimethylsilyl)propionic-2,2,3,3-d_4_ acid sodium salt (TSP)] was used to dilute samples. An aliquot (560 μl) of defrosted (on ice) spent medium was added to 140 μl of the NMR buffer, and the mixture was vortexed for 30 s before 600 μl was transferred to an NMR tube (Wilmad® NMR tubes 5 mm diam., economy) and stored for no more than 24 h at 4 °C before analysis on a Jeol ECX 400 instrument (^1^H: 400 MHz; NOESY with water pre-saturation). Each spectrum was acquired with 128 scans, using a relaxation delay of 5 s (1.1 s acquisition time) and processed with Jeol Delta 5.3 software. Resulting one-dimensional spectra were imported into Chenomx NMR Suite 10.1 (Chenomx Inc.), where they were batch-processed (phase correction, baseline correction – Whittaker spline, water region deleted, pH 7.4 ± 0.50, TSP 1 mM). Each spectrum was profiled manually with reference to Chenomx’s reference metabolite library. Manual profiling of metabolites was done, particularly for samples including AU, due to low complexity of spectra. Concentrations of metabolites detected in samples were divided by 0.8 to account for dilution with NMR buffer.

### Statistical analysis

Two-way ANOVA with interaction effect, with Tukey’s Honest Significant Differences test [with false discovery rate (Benjamini–Hochberg procedure) implemented], was used to analyse biofilm and NMR data (in terms of medium, atmosphere, medium:atmosphere) for each strain or sample dataset.

## RESULTS

In this study, we determined the ability of 29 strains of well-characterized CAUTI-associated Gram-negative bacteria [27] to produce biofilms in TSB, AU and HU under aerobic (21 % O_2_) or microaerobic (5 % O_2_) conditions. These strains represented five different species of bacteria across five genera: *Escherichia coli* (*n* = 8), *Klebsiella pneumoniae* (*n* = 4), *Pseudomonas aeruginosa* (*n* = 8), *Enterobacter hormaechei* (*n* = 4) and *Proteus mirabilis* (*n* = 5). TSB represented a standard laboratory medium, with AU and HU representing physiologically relevant media. Aerobic conditions represented the atmosphere in which routine work with uropathogens is done, whereas microaerobic conditions represented the hypoxic environment of the human bladder. We also used untargeted metabolomics (^1^H-NMR, 400 MHz) to characterize the metabolites produced by our strains when grown in AU and HU under the different atmospheric conditions. An overview of the approach used herein can be found in **Supplementary Figure 1**.

### Effects of growth medium and atmosphere on biofilm formation (crystal violet assay)

Under aerobic or microaerobic conditions, biofilm formation was generally higher in TSB (21/29 and 18/29 strains, respectively) than in AU or HU (8/29 and 11/29 strains, respectively), but this was strain-specific across species (**Figure 1**, **Supplementary Tables 1–3**).

**Figure 1.**
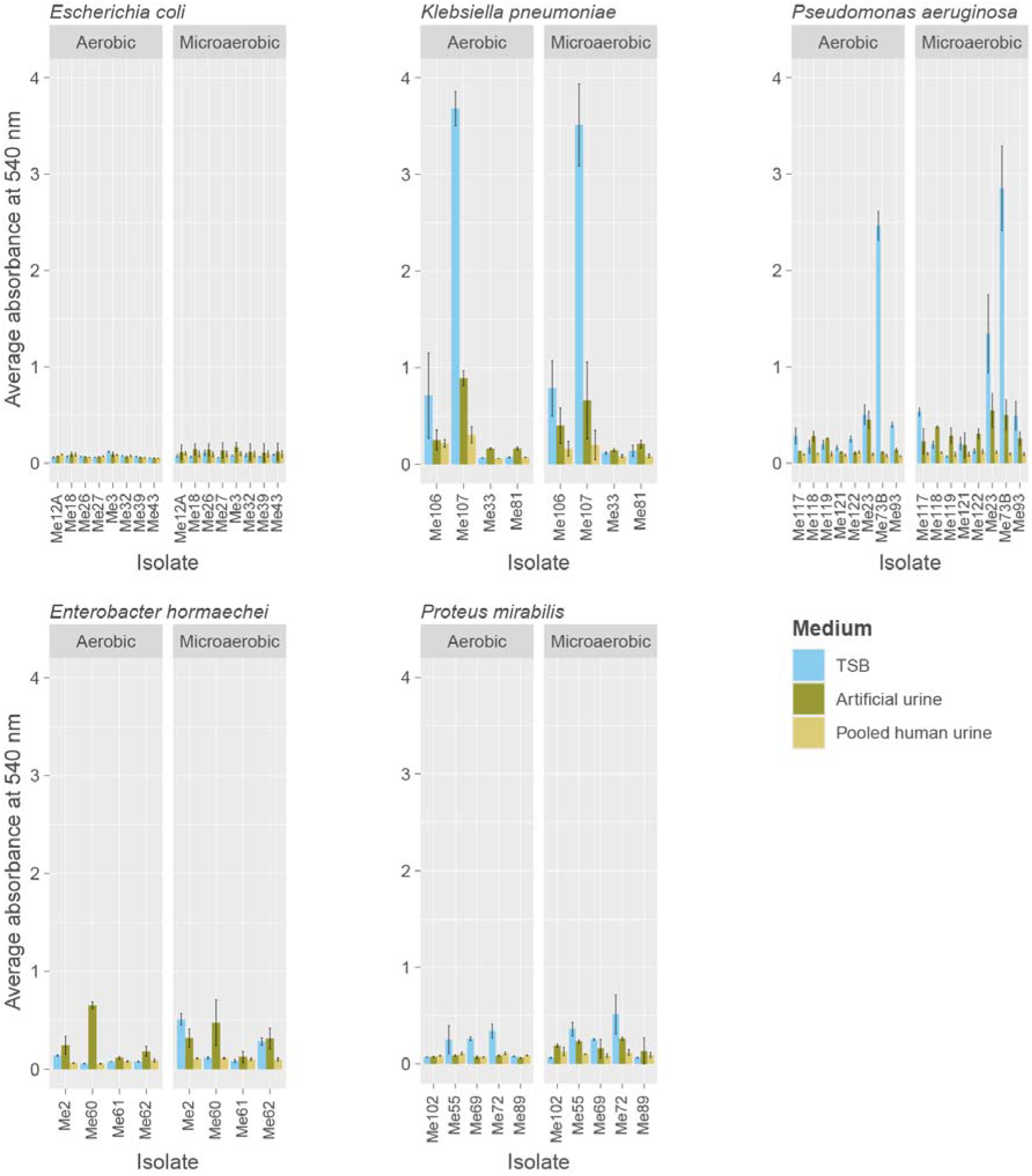
Assessment of medium (TSB, AU, HU) and atmosphere (aerobic, microaerobic) on biofilm formation by uropathogens. Data are shown as mean ± SD for three technical and three biological replicates per strain.

Mean biofilm formation by *Escherichia coli* strains was higher in HU and AU than in TSB under microaerobic conditions for 7/8 and 8/8 strains, respectively (**Supplementary Table 1**). Higher biofilm formation was seen in AU compared with HU for 5/8 and 8/8 strains, respectively, under aerobic and microaerobic conditions. Atmosphere significantly (P < 3.8 x 10^-2^) affected biofilm formation by strains Me26 and Me43, with a significant (P = 9.9 x 10^-3^) interaction effect (medium:atmosphere) observed for strain Me3 (**Supplementary Table 3**). Strain Me3 produced significantly (P < 2.97 x 10^-2^) more biofilm in AU when grown under microaerobic conditions compared with growth in AU and HU in aerobic conditions, and with growth in TSB in aerobic conditions. *Escherichia coli* strains tended to form the least biofilm across all growth condition and species combinations (**Figure 1**).

Growth medium significantly (P < 9.00 x 10^-3^) influenced biofilm formation by *Klebsiella pneumoniae*, with strain Me107 forming the most biofilm across all strains when incubated (micro)aerobically in TSB, AU or HU (**Figure 1**). Atmospheric conditions significantly (P < 3.53 x 10^-2^) influenced biofilm formation by strains Me107, Me33 and Me81 (**Supplementary Table 3**). A significant (P < .34 x 10^-2^) interaction effect (medium:atmosphere) was observed for the latter two strains, with significantly (P < 5.64 x 10^-3^) more biofilm formed by them when they were grown in the different media under microaerobic compared with aerobic conditions. Strain Me81 formed significantly (P = 1.26 x 10^-3^) more biofilm in AU than in TSB under aerobic conditions. This strain and Me33 formed significantly (P < 0.05) more biofilm in AU than in TSB under both atmospheric conditions. All four strains formed more biofilm in AU than in HU (P > 0.05 for Me106).

Atmosphere and growth medium significantly (P < 0.05) influenced biofilm formation by 6/8 and 7/8 *Pseudomonas aeruginosa* strains (**Supplementary Table 3**). More biofilm was formed in AU and TSB than HU for all strains (micro)aerobically (**Figure 1**, **Supplementary Table 1**). A significant (P < 0.05) interaction effect (medium:atmosphere) was seen for 3/8 strains (Me23, Me119 and Me122). Strain Me23 formed more biofilm in TSB and AU (micro)aerobically than in HU under both atmospheric conditions, forming significantly (P < 1.38 x 10^-2^) more biofilm in TSB incubated microaerobically than other growth conditions (**Figure 1**). Strain Me73B formed most biofilm in TSB among the *Pseudomonas aeruginosa* strains (micro)aerobically. Strains Me119 and Me122 formed significantly (P < 0.05) more biofilm in AU microaerobically than in HU (micro)aerobically or TSB (aerobically and non-significant, respectively).

Growth medium and atmosphere significantly influenced biofilm formation in 4/4 (P < 4.23 x 10^-2^) and 2/4 (P < 1.10 x 10^-3^) *Enterobacter hormaechei* strains, respectively, with the two strains (Me2, Me62) also subject to a significant (P < 2.47 x 10^-2^) interaction effect (medium:atmosphere) (**Supplementary Table 3**). Three of the four strains formed significantly (P < 0.05) more biofilm in AU than in HU (Me61 non-significant). Except for strain Me60 (AU, aerobic), the *Enterobacter hormaechei* strains formed most biofilm under microaerobic conditions across the three media (**Figure 1**, **Supplementary Table 1**).

As for the other four species, *Proteus mirabilis* strains were subject to significant (P < 0.05) strain-dependent medium and/or atmosphere effects (**Supplementary Table 3**). With the exceptions of strains Me102 and Me89 (TSB, aerobic), all strains formed most biofilm under microaerobic conditions across all media (**Figure 1**, **Supplementary Table 1**).

### Effects of growth medium and atmosphere on biofilm formation (cfu)

Although the crystal violet assay is used to assess biofilm formation, it does not allow quantification of the number of bacteria in biofilms. Consequently, we assessed the number of bacteria found in biofilms of a representative set of isolates after growth in TSB, AU and HU in (micro)aerobic conditions (**Figure 2**, **Supplementary Table 4**).

**Figure 2.**
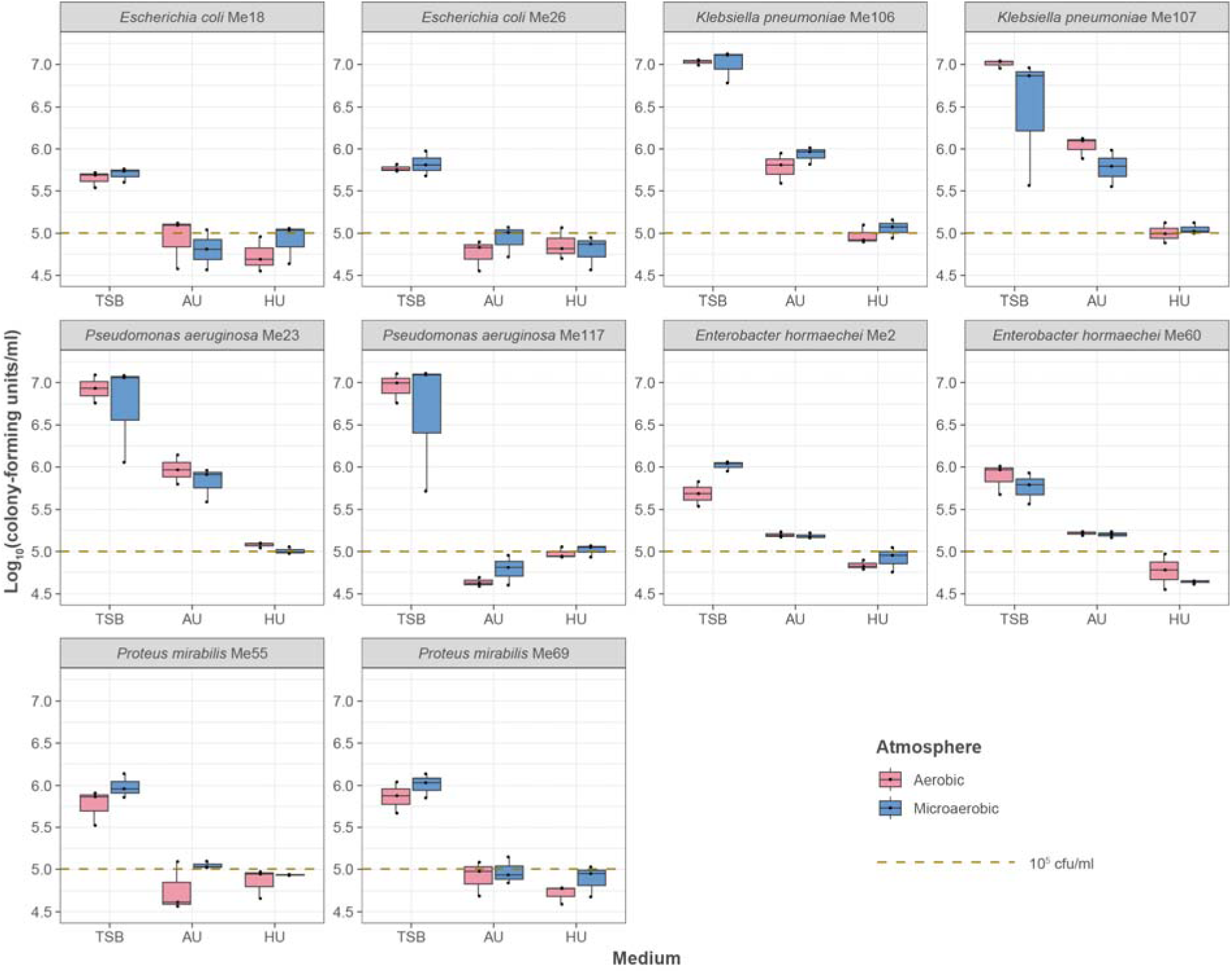
Representative data for effects of growth medium (AU, HU, TSB) and atmosphere (aerobic, microaerobic) on number of bacteria found in biofilms. Data were generated for two strains randomly selected for each species, and are shown for three biological replicates per strain.

Unsurprisingly, growth medium had a significant (P = 0) effect on bacteria recovered from biofilms formed by all strains (**Supplementary Table 5**). Across all strains (except Me2, TSB), no significant differences were found between microaerobic and aerobic biofilm formation within TSB, AU or HU (**Supplementary Table 5**). Atmosphere significantly (P = 3.46 x 10^-2^) affected number of bacteria recovered for *Proteus mirabilis* Me55 (it grew better upon microaerobic incubation in all media; **Figure 2**, **Supplementary Table 4**), while a significant (P = 9.80 x 10^-3^) interaction effect (medium:atmosphere) was noted for *Enterobacter hormaechei* Me2 [it grew significantly (P < 0.05) better in TSB and HU incubated microaerobically, and AU incubated aerobically]. Significantly (P < 0.05) more bacterial cells were recovered from biofilms grown in TSB than in AU or HU under aerobic and microaerobic conditions for all strains, except for *Klebsiella pneumoniae* Me107 (TSB, microaerobic; non-significance due to spread of data) (**Figure 2**, **Supplementary Table 5**). Biofilm formation in TSB was not comparable to that in the two physiologically relevant growth substrates (**Figure 2**). No further work was done with TSB samples.

Comparing the biofilm data generated for the crystal violet assay with the cfu data, the *Escherichia coli* strains produced fewest cells in the three media (**Figure 3**) in agreement with their low biofilm formation (**Figure 2**). *Escherichia coli* Me18 and Me26 formed more biofilm in AU incubated aerobically (**Figure 2**; **Supplementary Table 1**), but their cfu data showed TSB to be the medium producing most cells in biofilms (**Figure 3**; **Supplementary Table 4**). Similar results were seen for *Enterobacter hormaechei* Me60. Therefore, amount of biofilm formed determined using the crystal violet assay is not necessarily contingent upon the number of bacterial cells present in the biofilm.

**Figure 3.**
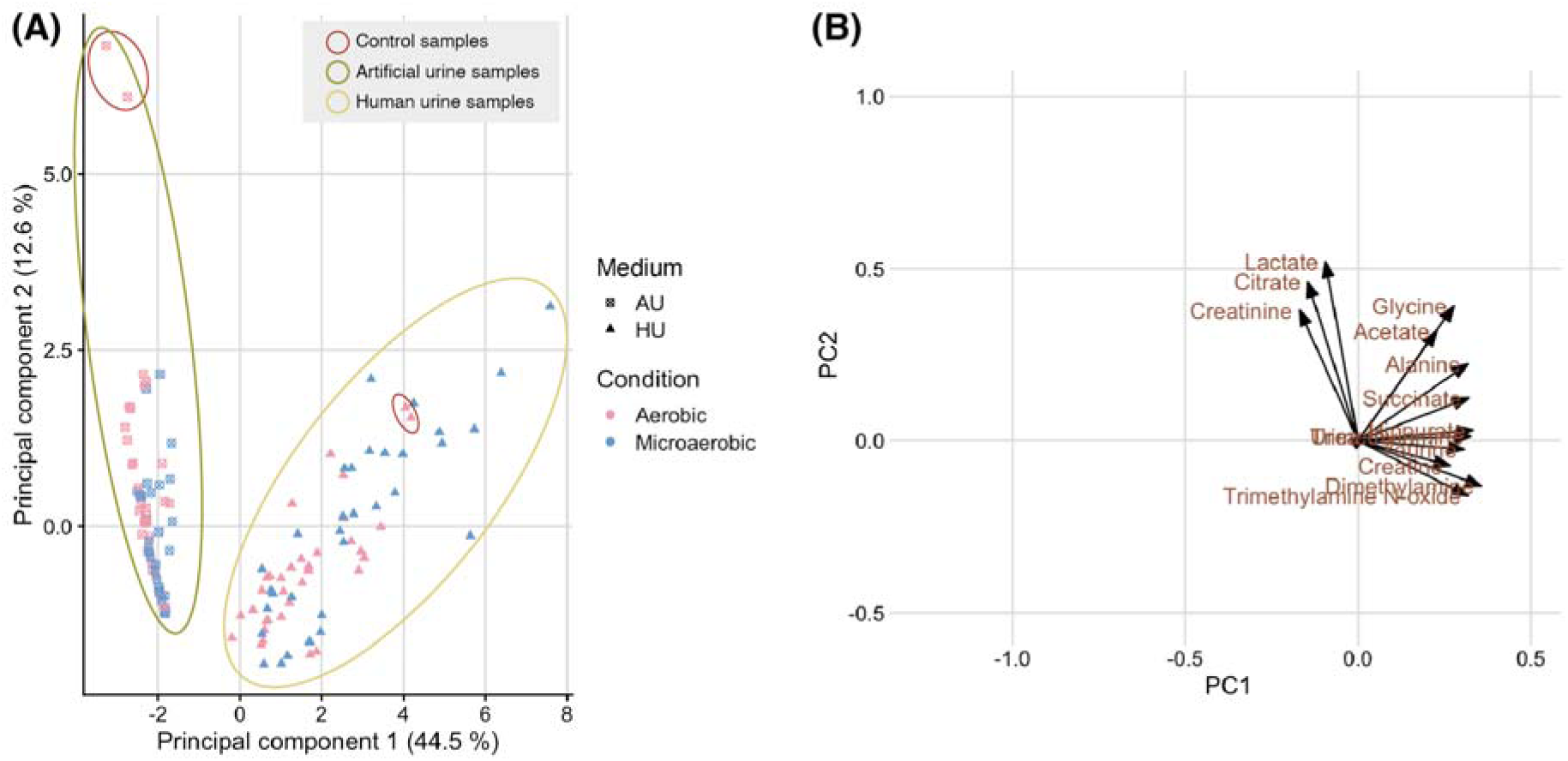
PCA of NMR data generated for the 14 metabolites quantified in this study. Mean, scaled data (*n* = 3 biological replicates) for 29 strains plus the negative (uninoculated) controls were used to generate the **(A)** scores and **(B)** loadings plots for the metabolomic data. The controls for AU and HU cluster together within their respective media clusters [circled in red in **(A)**]. *Escherichia coli*, *n* = 8; *Klebsiella pneumoniae*, *n* = 4; *Pseudomonas aeruginosa*, *n* = 8; *Enterobacter hormaechei*, *n* = 4; *Proteus mirabilis*, *n* = 5.

There was good agreement between the crystal violet data for the *Klebsiella pneumoniae*, *Pseudomonas aeruginosa* and *Proteus mirabilis* strains and the number of bacteria in biofilms across all three media and oxygen conditions (**Figure 2**, **Figure 3**). The (micro)aerobic data for *Pseudomonas aeruginosa* Me117 did not agree with respect to HU and AU – in the crystal violet assay more biofilm was produced in AU, but in HU more bacteria were found in the biofilm (**Supplementary Table 1**, **Supplementary Table 4**).

### The NMR-acquired metabolome of Gram-negative uropathogens

We wanted to determine how comparable AU was to HU with respect to its metabolite composition and how this influenced the metabolomes of uropathogens grown under aerobic and microaerobic conditions. NMR analyses of uninoculated AU and HU showed that – except for acetate – AU did not contain any detectable microbiome-associated metabolites (**Supplementary Figure 2**). The mean concentrations of creatinine, lactate and citrate present in AU were far higher than in HU: 973 μM vs 550 μM; 1743 μM vs 180 μM; and 2145 μM vs 363 μM, respectively. Alanine (165 μM), creatine (68 μM), DMA (83 μM), hippurate (126 μM), succinate (35 μM), taurine (202 μM), TMA (6 μM) and TMAO (71 μM) were present only in HU. There were broadly comparable amounts of urea (1248 μM vs 1022 μM), acetate (98 μM vs 92 μM) and glycine (182 vs 270 μM) present in the AU and HU, respectively.

The metabolomic profiles of spent media collected from the biofilms were analysed by NMR for all 29 strains. Principal component analysis (PCA) showed the data could be separated into two clusters – one for AU and one for HU (**Figure 3A**). Principal component 1 was responsible for 44.5 % of the variance in the data, with the metabolites lactate, citrate and creatinine driving separation of the data (**Figure 3B**). This is unsurprising given these metabolites were the most abundant in the AU used in this study.

No alanine, creatine, DMA, hippurate, succinate, taurine, TMA or TMAO was detected in any of the spent AU samples (**Figure 4**, **Supplementary Table 6**). The microbiome-associated metabolites acetate, hippurate, DMA, TMAO and TMA were detected in spent HU samples (**Figure 4**).

**Figure 4.**
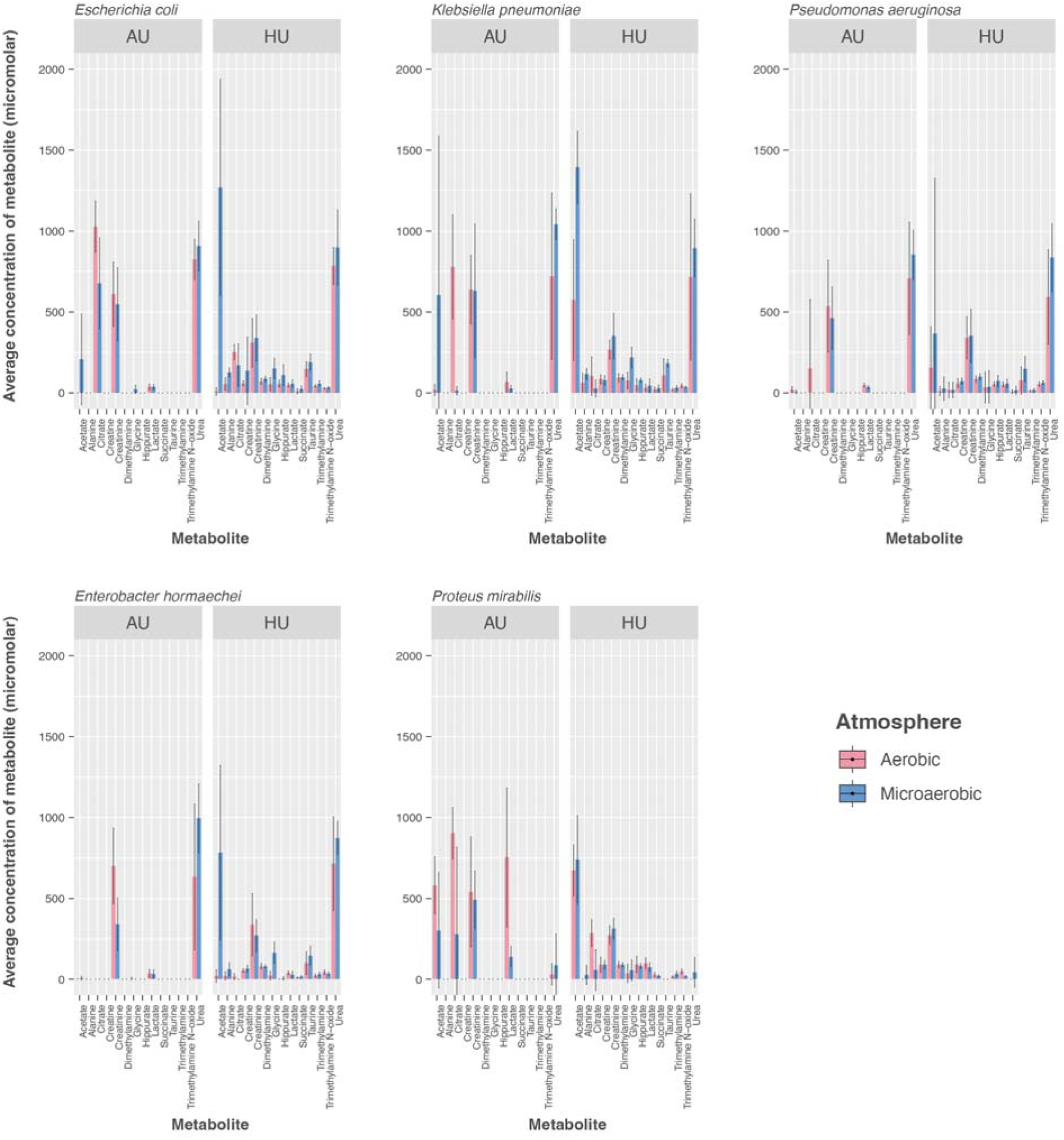
Summary of NMR data generated for 14 metabolites detected in this study. Aggregated data are presented as mean ± SD for three biological replicates for each strain of each species. *Escherichia coli*, *n* = 8; *Klebsiella pneumoniae*, *n* = 4; *Pseudomonas aeruginosa*, *n* = 8; *Enterobacter hormaechei*, *n* = 4; *Proteus mirabilis*, *n* = 5.

*Escherichia coli* produced significantly (P = 0.031) more acetate in microaerobic AU than aerobic AU, but this production was restricted to just three strains: Me12A, Me18 and M32 (**Figure 5A**, **Supplementary Table 7**). Three strains of *Klebsiella pneumoniae* (Me81, Me106, Me107) produced acetate from AU under microaerobic conditions, with one strain (Me106) producing creatinine. *Pseudomonas aeruginosa* and *Enterobacter hormaechei* strains were net consumers of all metabolites detected in spent AU under (micro)aerobic conditions, except for *Pseudomonas aeruginosa* Me93, which produced creatinine under aerobic conditions (**Supplementary Table 7**). Significantly more citrate was present in spent aerobic AU than spent microaerobic AU for *Escherichia coli* (P = 1.10 x 10^-7^), *Klebsiella pneumoniae* (P = 0.0046) and *Proteus mirabilis* (P = 3.83 x 10^-5^). Significantly (P = 0.0030) more creatinine and significantly (P = 0.0029) less urea were present in spent aerobic AU than spent microaerobic AU for *Enterobacter hormaechei*. All *Proteus mirabilis* strains produced acetate from aerobic AU, with Me102, Me69 and Me72 producing acetate from microaerobic AU. One strain (Me55) produced creatinine in aerobic AU.

**Figure 5.**
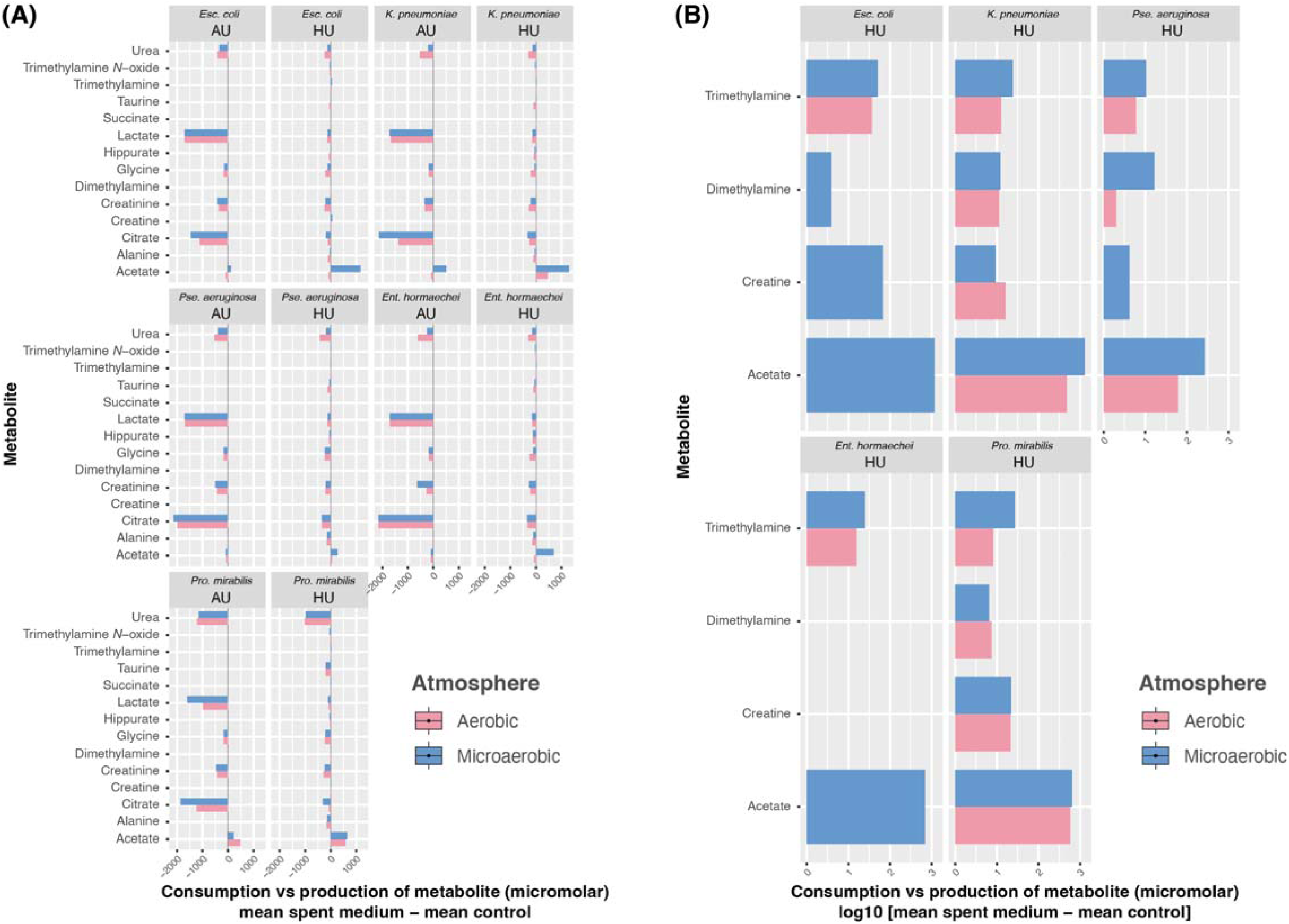
Consumption or production of metabolites in AU and HU, as determined by comparison to the uninoculated negative controls. Data in **(A)** and **(B)** are presented as the mean for each species: *Escherichia coli*, *n* = 8; *Klebsiella pneumoniae*, *n* = 4; *Pseudomonas aeruginosa*, *n* = 8; *Enterobacter hormaechei*, *n* = 4; *Proteus mirabilis*, *n* = 5. Data are log_10_-transformed in **(B)** to aid visualization of data for low-abundance metabolites.

*Escherichia coli* strains were net consumers of alanine, citrate, creatinine, glycine, hippurate, lactate, succinate and TMAO in HU (**Figure 5A**, **Supplementary Table 7**). A significant (P < 0.05) interaction between medium and atmosphere was observed for *Escherichia coli* (*n* = 8 strains) (**Supplementary Table 8**). Significant (P < 0.05) interactions were seen between metabolites and atmosphere for *Escherichia coli*, *Klebsiella pneumoniae* (*n* = 4 strains), *Enterobacter hormaechei* (*n* = 4 strains) and *Proteus mirabilis* (*n* = 5 strains); metabolite:interactions were significant for all five species (**Supplementary Table 8**).

All *Escherichia coli* strains produced significant (P = 7.57 x 10^-14^) quantities of acetate from HU under microaerobic compared with aerobic conditions (**Figure 4**, **Figure 5**). They were net consumers of alanine, citrate, lactate and TMAO, and net producers of TMA in both atmospheric conditions. Strains Me12 and Me43 produced creatine (micro)aerobically, with much more produced under microaerobic conditions by both strains (plus 9 μM vs 53 μM and 10 μM vs 574 μM compared with the negative control, respectively); creatine was also produced at low levels (plus 15 μM and 8 μM, respectively, compared with the negative control) microaerobically by strains Me18 and Me26. Strain Me27 produced creatinine (micro)aerobically. DMA was produced aerobically by strains Me3 and Me39, and microaerobically by strains Me12A, Me26, Me32 and Me39.

Except for strain Me107 (aerobic), all *Klebsiella pneumoniae* strains produced significantly (P = 8.75 x 10^-10^) more acetate from microaerobic HU than aerobic HU (**Figure 4**). They were all net consumers of alanine, citrate, creatinine, hippurate, lactate and TMAO, and net producers of TMA under (micro)aerobic conditions (**Figure 5**). Strains Me106 and Me107 produced creatine (micro)aerobically, while these strains and Me33 produced DMA (micro)aerobically.

Acetate was produced by *Enterobacter hormaechei* strains Me2, Me60 and Me62 under microaerobic conditions in HU (P = 3.74 x 10^-10^ when aerobic and microaerobic data compared); strains Me2 and Me62 produced creatinine microaerobically; strains Me2 (aerobic) and Me60 [(micro)aerobic] produced DMA; taurine was produced by strain Me2 under aerobic conditions, with strain Me60 producing urea aerobically.

All *Proteus mirabilis* strains produced acetate (micro)aerobically from HU. They were net consumers of alanine, creatinine [significantly (P = 4.10 x 10^-2^) more present in microaerobic spent HU], glycine, lactate, taurine, TMAO and urea (micro)aerobically. DMA was produced by strains Me102 and Me69 (micro)aerobically. Except for Me102 (aerobic), all *Proteus mirabilis* strains produced TMA; except for Me89 (aerobic), they were consumers of citrate, with significantly (P = 1.30 x 10^-3^) more present in spent aerobic HU than microaerobic HU.

*Pseudomonas aeruginosa* strains were net consumers of citrate and lactate. There was variability in the ability of the strains to produce acetate, with only two strains (Me117, Me118) able to produce this metabolite in HU under both aerobic and microaerobic conditions. Similarly, creatine and DMA production were variable. TMA was produced aerobically and microaerobically by 7/8 strains (except Me122 aerobically and Me93 microaerobically).

## DISCUSSION

The main aims of this study were to determine the effects of physiologically relevant media (AU and HU) and O_2_ availability on biofilm formation and the NMR-acquired metabolome of CAUTI-associated bacteria. A secondary aim was to see whether we could produce NMR metabolomic profiles akin to those seen in patients affected by UTIs [14–18], with a long-term goal of defining more appropriate growth conditions for uropathogens to better understand their physiology and host–microbiota interactions in the urinary tract.

Generating biofilm and metabolomic data for five of the species most often associated with CAUTIs – *Escherichia coli*, *Klebsiella pneumoniae*, *Pseudomonas aeruginosa*, *Enterobacter hormaechei* and *Proteus mirabilis* – we have shown that growth in HU leads to production of biofilms that often do not exceed 10^5^ cfu/ml, under either aerobic or microaerobic conditions. Bacterial counts recovered from midstream urine range from 10^3^ to <10^5^ cfu/ml in some patients with symptomatic UTIs, with the threshold of ≥10^5^ cfu/ml routinely used for diagnosis of UTIs [6,7]. We have also shown that growth of *Escherichia coli*, *Klebsiella pneumoniae* and *Proteus mirabilis* in HU under microaerobic conditions produces significantly higher quantities of acetate for all species, and small quantities of TMA, in agreement with NMR profiles obtained from urine of patients with UTIs caused by these Gram-negative bacteria [14–18].

For *Escherichia coli* and *Enterobacter hormaechei* we observed that biofilm formation assessed using the crystal violet assay does not reflect the number of bacterial cells present in the biofilm. This has important implications for how we assess bacterial load in biofilms formed by uropathogens, especially when comparing biofilm formation across different media and O_2_ concentrations. Multiple methods used in concert (including live/dead staining) are needed to assess biofilm formation, while acknowledging their inherent limitations in the interpretation of results [29].

Eberly et al. [30] found biofilm formation by UPEC decreased in a step-wise fashion from 21 % to 10 % oxygen when grown in AU, with biofilm levels at 4 % O_2_ the same as those seen under 21 % O_2_. Biofilm formation in their AU (comprising diluted TSB, oxalate, divalent cations, salts, citrate, ammonium, urea and creatinine) was higher overall by crystal violet staining compared to the levels of biofilm formed when grown in lysogeny broth [30]. We found medium composition (AU and HU specifically) had a greater effect on biofilm formation by UPEC, and saw higher biofilm formation microaerobically than aerobically.

To the best of our knowledge, this is the first study to characterize the NMR-acquired metabolome of CAUTI-associated bacteria using different growth conditions. We found that AU and HU do not produce comparable data. The composition of self-made [31–34] or commercially available AU needs to include microbiome-associated metabolites (e.g. hippurate, TMAO, DMA, TMA) routinely detected by NMR in HU if it is to be considered a true substitute to working with human biofluids. Eberly et al. [30] assessed the effect of TMAO, an alternative electron acceptor, on biofilm formation by UPEC under anaerobic conditions. Using supraphysiological concentrations (≥20 mM) they found no effect of TMAO on biofilm formation. TMAO is known to specifically encourage growth of *Enterobacteriaceae* under anaerobic conditions, resulting in the production of TMA [24]. It was surprising to us that TMA was produced from TMAO under (micro)aerobic conditions. Understanding how TMA production is influenced by O_2_ availability will be the subject of future work. In addition, we noted strain- and species-specific differences in the production of DMA, creatine and acetate from HU under (micro)aerobic conditions. As such, fundamental work is required to better understand the metabolic processes of uropathogenic bacteria under physiologically relevant conditions.

We acknowledge this proof-of-principle study has limitations. Chief among which is the relatively high limit of detection (∼1 µM) of NMR compared with methods such as liquid chromatography–mass spectrometry analysis [35]. Quantification of low-abundance metabolites such as TMA and creatine is thus semi-quantitative. We were also unable to detect by NMR many of the biogenic amines known to be produced in HU by Gram-negative uropathogens [35]. We found working with NMR data from AU-produced samples to be problematic: if bacteria used all components of the medium, we could not produce usable NMR spectra, hence why data are presented for only the 29 strains for which we had complete sets of AU and HU spectra. Single batches of AU and pooled HU were used in this study. Future work should consider sourcing urine from a range of individuals, including those with conditions such as diabetes, as glucose availability, age, sex and medications will affect the urinary metabolome and thereby growth of bacteria. Precipitate formation in AU was an issue in the crystal violet assay, but we do not think this influenced absorbance readings returned for biofilm formation by strains. We grew strains for 24 h to produce the biofilm and NMR data presented herein. This may be too long for Gram-negative bacteria. Indeed, Chan et al. [35] used a shorter time frame to characterize the metabolome of a range of aerobically grown Gram-negative (*Escherichia coli*, *Klebsiella pneumoniae*, *Pseudomonas aeruginosa*, *Proteus mirabilis*) and Gram-positive (*Enterococcus faecalis*, *Streptococcus agalactiae*, *Staphylococcus aureus*, *Staphylococcus saprophyticus*) uropathogens, and were able to demonstrate the different bacterial species could be assigned to groups based on serine consumption (*Escherichia coli*, *Klebsiella pneumoniae*, *Proteus mirabilis*), glutamine consumption (*Pseudomonas aeruginosa*), amino acid abstention (*Streptococcus agalactiae*, *Enterococcus faecali*s) and amino acid minimalism (staphylococci).

## ACKNOWLEDGEMENTS

Computing resources used in this study were funded through the Research Contingency Fund of Nottingham Trent University.

## AUTHOR CONTRIBUTIONS

ME undertook all phenotypic work. CJG, ALM and LH did all NMR work. LH annotated and analysed all NMR data. LH, ME and ALM wrote the manuscript. ME and LH secured funding for the work. All authors approved the final version of the manuscript.

## Abbreviations

AU: artificial urine
CAUTI: catheter-associated urinary tract infection
cfu: colony-forming unit
DMA: dimethylamine
HU: human urine
NMR: ^1^H-nuclear magnetic resonance spectroscopy
PCA: principal component analysis
PBS: phosphate-buffered saline
TMA: trimethylamine
TMAO: trimethylamine *N*-oxide
TSB: tryptone soya broth containing 1 % glucose
UPEC: uropathogenic *Escherichia coli*
UTI: urinary tract infection

---

**Supplementary Figure 1.**
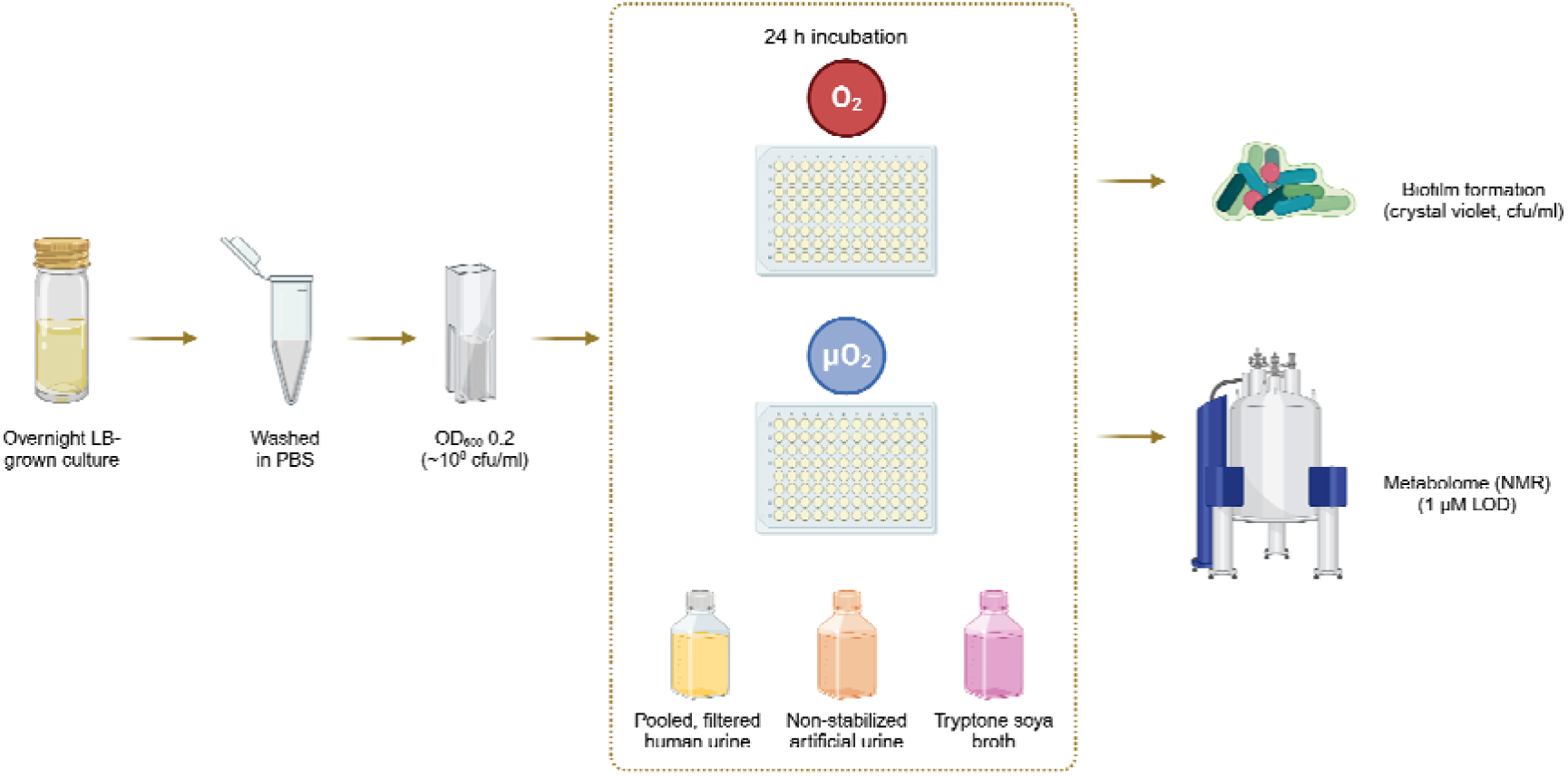
Overview of the simple methodological approach used in this study. Uropathogens were cultivated in broth overnight. Cell pellets were then washed to remove spent media and adjusted to a standard optical density in PBS, before being used to inoculate plates containing HU, AU or TSB. We left the plates to incubate aerobically or microaerobically for 24 h, then assessed biofilm formation and metabolomic profiles. LOD, limit of detection. Created in BioRender. Hoyles, L. (2025) https://BioRender.com/oqepb13.

**Supplementary Figure 2.**
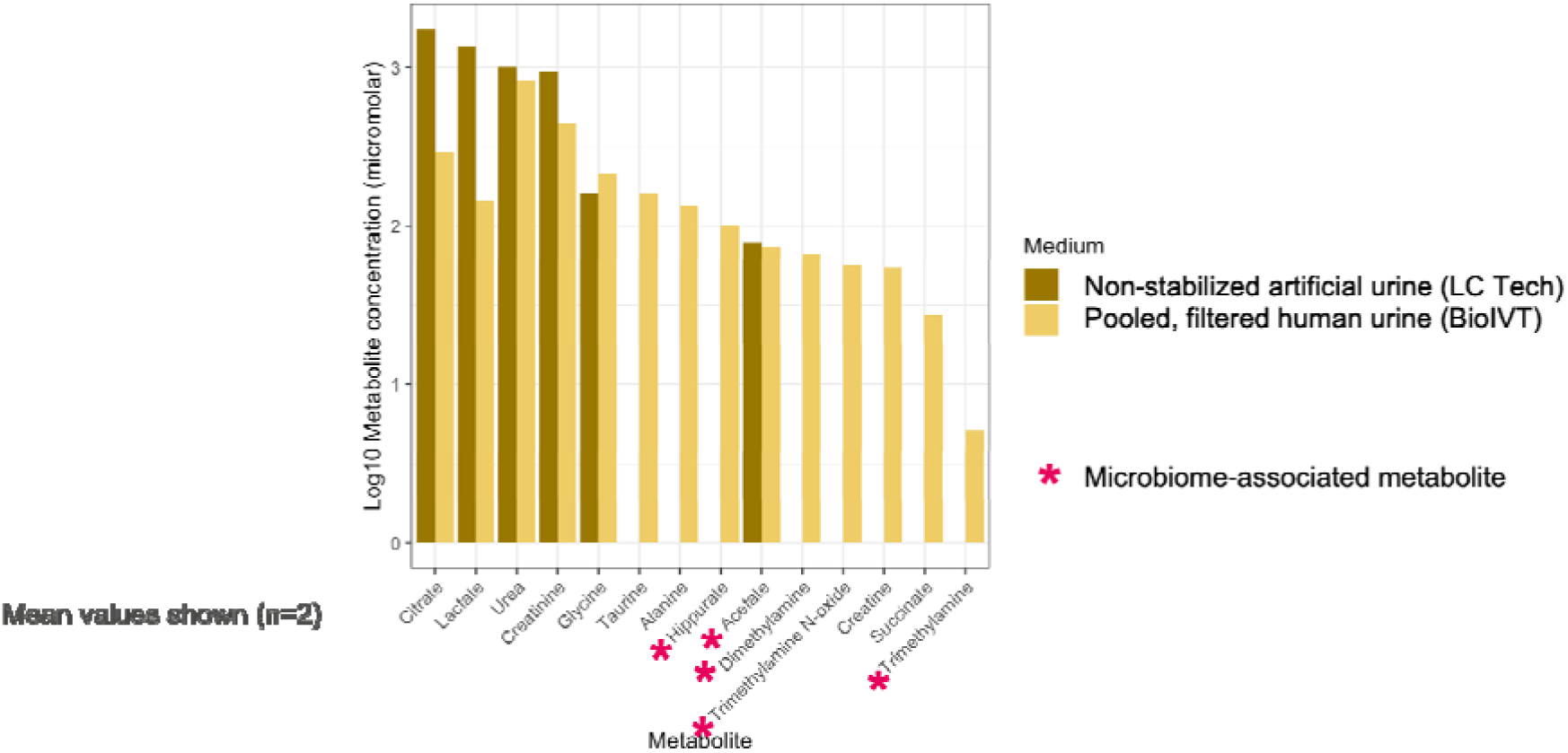
Composition of AU and HU used in this study as determined by ChenoMx analysis of NMR spectra. Data are shown as mean of two independent measurements. Metabolite concentrations were determined using ChenoMx Pro with reference to the TSP internal standard (1 μM final concentration).

## REFERENCES

[1] Advani SD, Thaden JT, Perez R, Stair SL, Lee UJ, Siddiqui NY. State-of-the-Art Review: Recurrent Uncomplicated Urinary Tract Infections in Women. Clin Infect Dis Off Publ Infect Dis Soc Am 2025;80:e31–42. 10.1093/cid/ciae653.

[2] Mancuso G, Midiri A, Gerace E, Marra M, Zummo S, Biondo C. Urinary Tract Infections: The Current Scenario and Future Prospects. Pathog Basel Switz 2023;12:623. 10.3390/pathogens12040623.

[3] Haque M, Sartelli M, McKimm J, Abu Bakar M. Health care-associated infections - an overview. Infect Drug Resist 2018;11:2321–33. 10.2147/IDR.S177247.

[4] Pothoven R. Management of urinary tract infections in the era of antimicrobial resistance. Drug Target Insights 2023;17:126–37. 10.33393/dti.2023.2660.

[5] McCann E, Sung AH, Ye G, Vankeepuram L, Tabak YP. Contributing Factors to the Clinical and Economic Burden of Patients with Laboratory-Confirmed Carbapenem-Nonsusceptible Gram-Negative Urinary Tract Infections. Clin Outcomes Res CEOR 2020;12:191–200. 10.2147/CEOR.S234840.

[6] Schmiemann G, Kniehl E, Gebhardt K, Matejczyk MM, Hummers-Pradier E. The diagnosis of urinary tract infection: a systematic review. Dtsch Arzteblatt Int 2010;107:361–7. 10.3238/arztebl.2010.0361.

[7] Little P, Turner S, Rumsby K, Warner G, Moore M, Lowes JA, et al. Developing clinical rules to predict urinary tract infection in primary care settings: sensitivity and specificity of near patient tests (dipsticks) and clinical scores. Br J Gen Pract J R Coll Gen Pract 2006;56:606–12.

[8] Public Health England. Exploring the implementation of interventions to reduce catheter-associated urinary tract infections (ENACT). 2021.

[9] Shannon MB, Limeira R, Johansen D, Gao X, Lin H, Dong Q, et al. Bladder urinary oxygen tension is correlated with urinary microbiota composition. Int Urogynecology J 2019;30:1261–7. 10.1007/s00192-019-03931-y.

[10] Hu RT, Lankadeva YR, Yanase F, Osawa EA, Evans RG, Bellomo R. Continuous bladder urinary oxygen tension as a new tool to monitor medullary oxygenation in the critically ill. Crit Care Lond Engl 2022;26:389. 10.1186/s13054-022-04230-7.

[11] Wang ZJ, Joe BN, Coakley FV, Zaharchuk G, Busse R, Yeh BM. Urinary oxygen tension measurement in humans using magnetic resonance imaging. Acad Radiol 2008;15:1467–73. 10.1016/j.acra.2008.04.013.

[12] Giannakopoulos X, Evangelou A, Kalfakakou V, Grammeniatis E, Papandropoulos I, Charalambopoulos K. Human bladder urine oxygen content: implications for urinary tract diseases. Int Urol Nephrol 1997;29:393–401. 10.1007/BF02551103.

[13] Abosamak MF, Lippi G, Benoit SW, Henry BM, Shama AAA. Bladder urine oxygen partial pressure monitoring: Could it be a tool for early detection of acute kidney injury? Egypt J Anaesth 2021.

[14] Gupta A, Dwivedi M, Mahdi AA, Khetrapal CL, Bhandari M. Broad identification of bacterial type in urinary tract infection using (1)h NMR spectroscopy. J Proteome Res 2012;11:1844–54. 10.1021/pr2010692.

[15] Nevedomskaya E, Pacchiarotta T, Artemov A, Meissner A, van Nieuwkoop C, van Dissel JT, et al. (1)H NMR-based metabolic profiling of urinary tract infection: combining multiple statistical models and clinical data. Metabolomics Off J Metabolomic Soc 2012;8:1227–35. 10.1007/s11306-012-0411-y.

[16] Lam C-W, Law C-Y, To KK-W, Cheung SK-K, Lee K-C, Sze K-H, et al. NMR-based metabolomic urinalysis: a rapid screening test for urinary tract infection. Clin Chim Acta Int J Clin Chem 2014;436:217–23. 10.1016/j.cca.2014.05.014.

[17] Lam C-W, Law C-Y, Sze K-H, To KK-W. Quantitative metabolomics of urine for rapid etiological diagnosis of urinary tract infection: evaluation of a microbial-mammalian co-metabolite as a diagnostic biomarker. Clin Chim Acta Int J Clin Chem 2015;438:24–8. 10.1016/j.cca.2014.07.038.

[18] Lussu M, Camboni T, Piras C, Serra C, Del Carratore F, Griffin J, et al. 1H NMR spectroscopy-based metabolomics analysis for the diagnosis of symptomatic E. coli-associated urinary tract infection (UTI). BMC Microbiol 2017;17:201. 10.1186/s12866-017-1108-1.

[19] Russell WR, Hoyles L, Flint HJ, Dumas M-E. Colonic bacterial metabolites and human health. Curr Opin Microbiol 2013;16:246–54. 10.1016/j.mib.2013.07.002.

[20] Hoyles L, Pontifex MG, Rodriguez-Ramiro I, Anis-Alavi MA, Jelane KS, Snelling T, et al. Regulation of blood-brain barrier integrity by microbiome-associated methylamines and cognition by trimethylamine N-oxide. Microbiome 2021;9:235. 10.1186/s40168-021-01181-z.

[21] Nakamura J, Mutlu E, Sharma V, Collins L, Bodnar W, Yu R, et al. The endogenous exposome. DNA Repair 2014;19:3–13. 10.1016/j.dnarep.2014.03.031.

[22] Bouatra S, Aziat F, Mandal R, Guo AC, Wilson MR, Knox C, et al. The human urine metabolome. PloS One 2013;8:e73076. 10.1371/journal.pone.0073076.

[23] Brial F, Chilloux J, Nielsen T, Vieira-Silva S, Falony G, Andrikopoulos P, et al. Human and preclinical studies of the host–gut microbiome co-metabolite hippurate as a marker and mediator of metabolic health. Gut 2021:gutjnl-2020–323314. 10.1136/gutjnl-2020-323314.

[24] Hoyles L, Jiménez-Pranteda ML, Chilloux J, Brial F, Myridakis A, Aranias T, et al. Metabolic retroconversion of trimethylamine N-oxide and the gut microbiota. Microbiome 2018;6:73. 10.1186/s40168-018-0461-0.

[25] Hoyles L, Fernández-Real J-M, Federici M, Serino M, Abbott J, Charpentier J, et al. Molecular phenomics and metagenomics of hepatic steatosis in non-diabetic obese women. Nat Med 2018;24:1070–80. 10.1038/s41591-018-0061-3.

[26] Koh A, Molinaro A, Ståhlman M, Khan MT, Schmidt C, Mannerås-Holm L, et al. Microbially Produced Imidazole Propionate Impairs Insulin Signaling through mTORC1. Cell 2018;175:947–961.e17. 10.1016/j.cell.2018.09.055.

[27] Eladawy M, Heslop N, Negus D, Thomas J, Hoyles L. Phenotype-genotype discordance in antimicrobial resistance profiles of Gram-negative uropathogens recovered from catheter-associated urinary tract infections in Egypt 2025. 10.1101/2025.04.17.649370.

[28] Eladawy M, Thomas JC, Hoyles L. Phenotypic and genomic characterization of Pseudomonas aeruginosa isolates recovered from catheter-associated urinary tract infections in an Egyptian hospital. Microb Genomics 2023;9. 10.1099/mgen.0.001125.

[29] Latka A, Leiman PG, Drulis-Kawa Z, Briers Y. Modeling the Architecture of Depolymerase-Containing Receptor Binding Proteins in Klebsiella Phages. Front Microbiol 2019;10:2649. 10.3389/fmicb.2019.02649.

[30] Eberly AR, Floyd KA, Beebout CJ, Colling SJ, Fitzgerald MJ, Stratton CW, et al. Biofilm Formation by Uropathogenic Escherichia coli Is Favored under Oxygen Conditions That Mimic the Bladder Environment. Int J Mol Sci 2017;18:2077. 10.3390/ijms18102077.

[31] Brooks T, Keevil CW. A simple artificial urine for the growth of urinary pathogens. Lett Appl Microbiol 1997;24:203–6. 10.1046/j.1472-765x.1997.00378.x.

[32] Chutipongtanate S, Thongboonkerd V. Systematic comparisons of artificial urine formulas for in vitro cellular study. Anal Biochem 2010;402:110–2. 10.1016/j.ab.2010.03.031.

[33] Sarigul N, Korkmaz F, Kurultak İ. A New Artificial Urine Protocol to Better Imitate Human Urine. Sci Rep 2019;9:20159. 10.1038/s41598-019-56693-4.

[34] Rimbi PT, O’Boyle N, Douce GR, Pizza M, Rosini R, Roe AJ. Enhancing a multi-purpose artificial urine for culture and gene expression studies of uropathogenic Escherichia coli strains. J Appl Microbiol 2024;135:lxae067. 10.1093/jambio/lxae067.

[35] Chan CCY, Groves RA, Rydzak T, Lewis IA. Metabolomics survey of uropathogenic bacteria in human urine. Front Microbiol 2024;15:1507561. 10.3389/fmicb.2024.1507561.

[36] Magiorakos A-P, Srinivasan A, Carey RB, Carmeli Y, Falagas ME, Giske CG, et al. Multidrug-resistant, extensively drug-resistant and pandrug-resistant bacteria: an international expert proposal for interim standard definitions for acquired resistance. Clin Microbiol Infect Off Publ Eur Soc Clin Microbiol Infect Dis 2012;18:268–81. 10.1111/j.1469-0691.2011.03570.x.

